# Novel genome-wide associations for anhedonia, genetic correlation with psychiatric disorders, and polygenic association with brain structure

**DOI:** 10.1101/656298

**Authors:** Joey Ward, Laura M. Lyall, Richard A. I. Bethlehem, Amy Ferguson, Rona J. Strawbridge, Donald M. Lyall, Breda Cullen, Nicholas Graham, Keira J.A. Johnston, Mark E.S. Bailey, Graham K. Murray, Daniel J. Smith

**Affiliations:** Institute of Health and Wellbeing, University of Glasgow, Glasgow, UK; Department of Psychiatry, University of Cambridge, Cambridge, UK; Department of Medicine Solna, Karolinska Institute, Stockholm, Sweden; School of Life Sciences, College of Medical, Veterinary and Life Sciences, University of Glasgow, Glasgow, UK

## Abstract

Anhedonia is a core feature of several psychiatric disorders but its biological underpinnings are poorly understood. We performed a genome-wide association study of anhedonia in 375,275 UK Biobank participants and assessed for genetic correlation between anhedonia and neuropsychiatric conditions (major depressive disorder, schizophrenia, bipolar disorder, obsessive compulsive disorder and Parkinson’s Disease). We then used a polygenic risk score approach to test for association between genetic loading for anhedonia and both brain structure and brain function. This included: magnetic resonance imaging (MRI) assessments of total grey matter volume, white matter volume, cerebrospinal fluid volume, and 15 cortical/subcortical regions of interest; diffusion tensor imaging (DTI) measures of white matter tract integrity; and functional MRI activity during an emotion processing task. We identified 11 novel loci associated at genome-wide significance with anhedonia, with a SNP heritability estimate (h_2_SNP) of 5.6%. Strong positive genetic correlations were found between anhedonia and major depressive disorder, schizophrenia and bipolar disorder; but not with obsessive compulsive disorder or Parkinson’s Disease. Polygenic risk for anhedonia was associated with poorer brain white matter integrity, smaller total grey matter volume, and smaller volumes of brain regions linked to reward and pleasure processing, including nucleus accumbens, caudate and medial frontal cortex. In summary, the identification of novel anhedonia-associated loci substantially expands our current understanding of the biological basis of anhedonia and genetic correlations with several psychiatric disorders confirm the utility of this trait as a transdiagnostic marker of vulnerability to mental illness. We also provide the first evidence that genetic risk for anhedonia influences brain structure, particularly in regions associated with reward and pleasure processing.

## Introduction

Anhedonia refers to reduced capacity to experience pleasure in situations that individuals would normally enjoy. It constitutes a core symptom of several neuropsychiatric disorders, particularly major depressive disorder (MDD), schizophrenia, bipolar disorder and obsessive compulsive disorder (OCD), as well as Parkinson’s Disease (PD)^1–4^. Along with a direct negative association with quality of life and subjective wellbeing, anhedonia is associated with multiple negative health-related behaviours, including smoking, illicit drug use, and lower physical activity, even in the absence of psychiatric disorder^5, 6^. In line with a Research Domain Criteria (RDoC) perspective^7^, anhedonia can be measured and studied as a dimensional psychopathological trait.

Anhedonia has been closely linked to the function and structure of reward circuitry in the brain (primarily frontal, striatal and limbic regions). These neurobiological associations are consistent with the view that anhedonia reflects dysfunction in reward processing^8, 9^. Measures of anhedonia have been associated with altered functional activity during reward-based tasks within frontal cortical regions (medial frontal cortex) and subcortical striatal regions (caudate and putamen)^10^, and with reduced volumes in a similar set of frontal and striatal regions^11, 12^. Anhedonia is also associated with reduced white matter integrity^13, 14^.

The genetic underpinnings of anhedonia are largely unknown. Several GWAS of disorders where anhedonia is a feature have been reported, such as MDD and schizophrenia^15, 16^. However, to date only small underpowered GWAS studies of anhedonia have been published. A study of 759 patients with MDD identified 18 SNPs associated with an ‘interest-activity’ measure of anhedonia^17^. The largest study to date is a mega-analysis of three studies of young people from the UK and Sweden, with a total sample size of 6579; a single locus was associated with anhedonia in the discovery sample, but not in the replication sample^18^. A Finnish study examined genetic associations with physical and social anhedonia, as assessed with the Chapman scales,^19^ in 3820 individuals but no genome-wide significant loci were identified^20^. Genetic loci associated with symptoms of anhedonia have therefore not yet been reliably identified in large clinical or general population samples^21^, and association tests of genetic risk for anhedonia with brain structure and function have not yet been performed.

Here we report a large GWAS of anhedonia within the UK Biobank cohort. We also use a polygenic risk score (PRS) approach to assess whether genetic loading for anhedonia is associated with brain structure and brain function.

## Methods

### UK Biobank sample

UK Biobank is a large cohort of over half a million UK residents, aged between 39 and 73 years at baseline assessment^22^. The cohort was designed to assess how genetic, environmental and lifestyle factors influence a range of morbidities in middle and older age. Baseline assessments occurred over a 4-year recruitment period (from 2006 to 2010) across 22 UK centres. These assessments covered a wide range of social, cognitive, lifestyle and physical health measures. Informed consent was obtained from all participants, and this study was conducted under generic approval from the NHS National Research Ethics Service (approval letter dated 13 May 2016, Ref 16/NW/0274) and under UK Biobank approvals for application #6553 ‘Genome-wide association studies of mental health’ (PI Smith).

### Genotyping, imputation and quality control

In March 2018 UK Biobank released genetic data for 487,409 individuals, genotyped using the Affymetrix UK BiLEVE Axiom or the Affymetrix UK Biobank Axiom arrays (Santa Clara, CA, USA), which have over 95% of content in common^23^. Pre-imputation quality control, imputation (both 1000Genomes and HRC Reference Panels) and post-imputation cleaning were conducted centrally by UK Biobank (described in the UK Biobank release documentation, please see URLs in appendix for details).

### Phenotyping

As part of the comprehensive baseline assessment participants were asked: *“Over the past two weeks, how often have you had little interest or pleasure in doing things?”* (Data field 2060). Respondents could choose from the following answers: *“not at all”; “several days”; “more than half the days”; and “nearly every day”.* These responses were coded as 0, 1, 2 and 3 respectively. This question on anhedonia is derived from the Patient Health Questionnaire-9 (PHQ-9), a well-validated screening instrument for MDD^24^. To maximise numbers available for downstream magnetic resonance imaging (MRI) analyses, we excluded from the primary GWAS those participants with any available MRI data. Additional exclusion criteria for the GWAS included individuals in whom: over 10% of genetic data were missing; self-reported sex did not match genetic sex; sex chromosome aneuploidy was reported; where heterozygosity value was a clear outlier; and participants not of European ancestry.

### Genetic association and heritability

Genetic association with the measure of anhedonia was performed using BOLT-LMM^25, 26^, which accounts for population structure and sample relatedness by including a genetic relatedness matrix within the models. Models were further adjusted for age, sex and genotyping array. SNPs included in the analysis were filtered by MAF > 0.01, Hardy-Weinberg Equilibrium p > 1×10^−6^, and imputation score > 0.3. BOLT-LMM was also used to provide a SNP-heritability estimate and an estimate of λ_GC_.

### Genetic correlations

Linkage Disequilibrium Score Regression (LDSR)^27^ was carried out using LDSC on GWAS summary statistics from several published studies, to obtain genetic correlations with psychiatric disorders where anhedonia is known to be a feature (MDD, schizophrenia, bipolar disorder and OCD), as well as Parkinson’s Disease.

### Polygenic risk score generation

Polygenic risk scores (PRS) were created using LDpred^28^. LDpred differs from the more common pruning and threshold (P&T) PRS because it generates a single risk score for the trait of interest derived from as many loci as possible. A training set was created to obtain the LD structure by using 1000 unrelated Biobank participants who had passed the same genetic QC as those used in the GWAS (but who were excluded from the GWAS because they did not respond to the anhedonia question and had not provided brain imaging data). These training data were then used for the construction of anhedonia PRS in those participants for whom brain imaging data were available.

### Brain imaging variables

A number of structural and functional brain MRI measures have been made available by UK Biobank as Imaging Derived Phenotypes (IDPs)^29^. These measures are: total volume (mm^3^) of brain grey matter, white matter and cerebrospinal fluid (CSF), each normalised for head size; volumes (grey matter or total) of 15 cortical and subcortical regions of interest (ROIs); diffusion tensor imaging (DTI) measures of white matter integrity (fractional anisotropy (FA) and mean diffusivity (MD)); and functional MRI activity during an emotion processing task (the Hariri face shape task)^30^ in an amygdala mask and group-defined mask consisting of occipito-temporal and amygdala regions. For a more detailed description of these variables, and for details of MRI acquisition and pre-processing and IDP selection, please see supplementary methods.

### Polygenic risk score and brain imaging analyses

MRI data were available for 20,174 UK Biobank participants. PRS/MRI analyses were conducted in a subset of 17,120 participants who had available MRI data and who were not included in the GWAS, after exclusion of participants who did not meet genetic quality control criteria (n = 2,479) or who self-reported a developmental or neurological disorder at either the baseline assessment or the imaging visit (n = 575) (please see Table S1 for exclusions). For each MRI outcome, data points with values more than 3 standard deviations from the sample mean were excluded.

Models were adjusted for age at MRI visit, (age at MRI)^2^, sex, genotype array, the first eight genetic principal components, and lateral, transverse and longitudinal scanner position covariates. Total tissue volume measures (total grey matter, white matter and ventricular CSF volumes) were normalised for head size prior to adjustment for the above covariates. ROI analyses were additionally adjusted for total brain volume (calculated by summing total grey matter, white matter and ventricular cerebrospinal fluid (CSF) volume); and fMRI analyses were also adjusted for head motion during the emotion processing task. False discovery rate (FDR) correction was applied^31, 32^.

## Results

### Demographics

The GWAS was performed on 375,724 UK Biobank participants, of whom 203,322 (54.1%) were female. The age of the sample ranged from 39 to 73 and the mean age was 57 years (S.D. = 8.01). In response to the anhedonia question, 299,232 (79.64%) answered *“not at all”*; 60,212 (16.0%) reported *“several days”*; 9,405 (2.5%) reported *“more than half the days”* and 6,876 (1.8%) reported *“nearly every day”.*

MRI analyses were conducted in 17,120 participants, 52.4% (8,978) of whom were female. The mean age (at the time of MRI) of these participants was 62.7 years (S.D. = 7.46; range = 45-80 years). Of the 16,783 participants with available MRI data who answered the anhedonia question, 13,810 (82.3%) responded *“not at all”,* 2,469 (14.7%) reported feelings of anhedonia for *“several days”,* 297 (1.8%) for *“more than half the days”,* and 207 (1.2%) *“nearly every day”.*

### Genome-wide association study findings

GWAS results are presented as a Manhattan plot in Figure 1, and details of genome-wide significant loci are provided in Table S2. In all, there were 1100 SNPs that were genome-wide significant (p < 5 x 10^−8^), and, of these, 11 represented independent lead SNPs at separate loci on 9 different chromosomes (Table S2; Figures S1-S11). An independent signal was defined as the region of r^2^ > 0.1 within a 500MB window from the most significant SNP below genome-wide significance.

**Figure 1.**
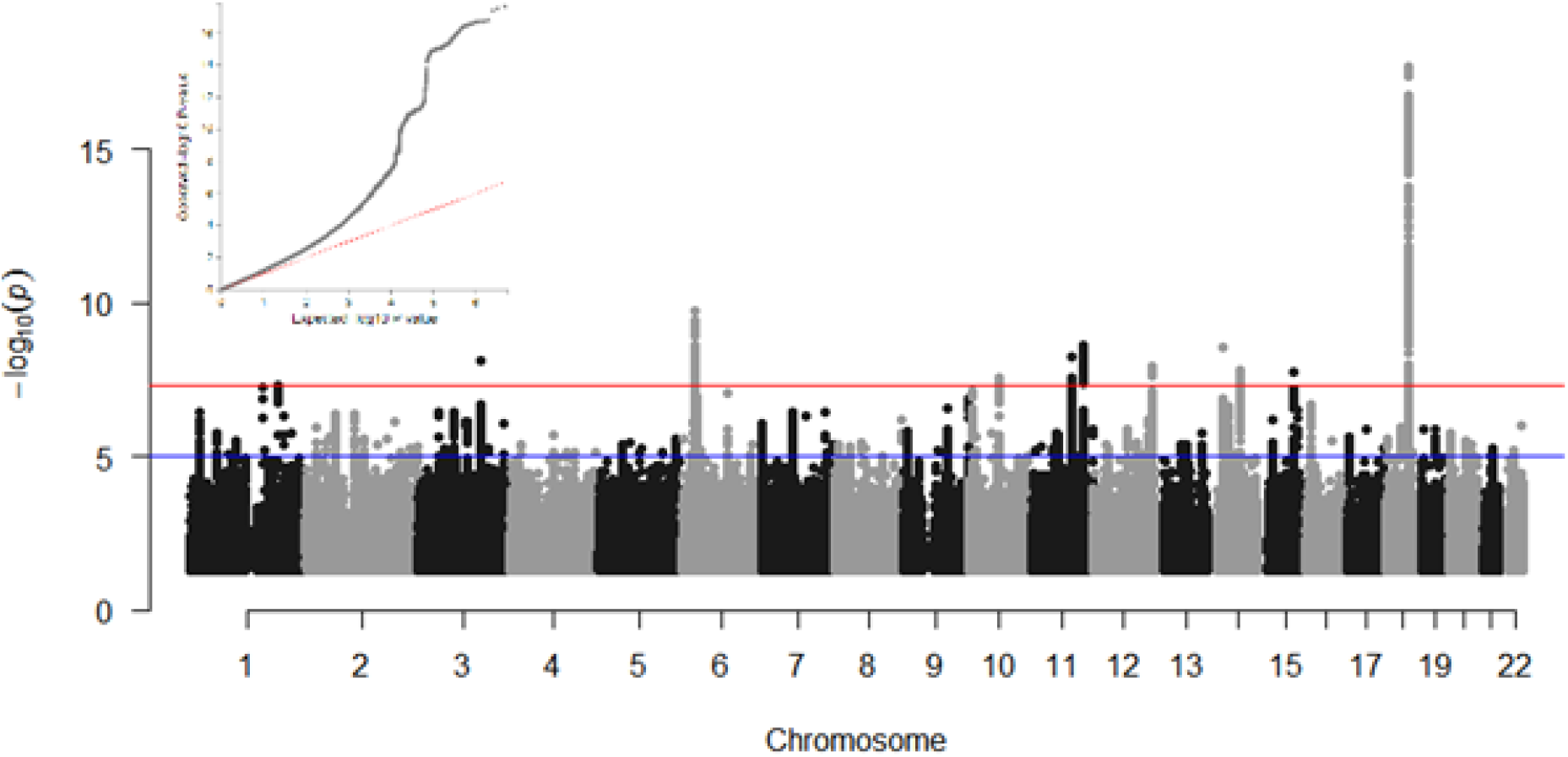
Anhedonia GWAS results. Results are presented as a Manhattan Plot and as a QQ plot (inset). Association analysis p-values for each SNP are plotted (as -log10(*p*)) vs. chromosomal position. The red and blue lines indicate the genome-wide significant and suggestive p-value thresholds, respectively. The QQ plot shows observed vs expected p-values for every SNP.

Some inflation of the GWAS results was observed (λ_GC_ = 1.15), however considering the sample size this is expected to have had a negligible impact on findings. There was evidence for a polygenic component (LDSR intercept = 1.03, S.E. = 0.005) and no evidence for undue inflation of the test statistics due to unaccounted population stratification.

### Genetic correlations

Using LDSR we assessed whether genetic predisposition to anhedonia overlapped with that for several psychiatric disorders or traits. Anhedonia had significant genetic correlation with MDD (r_g_ 0.77), schizophrenia (r_g_ 0.28) and bipolar disorder (r_g_ 0.12) but not with OCD or Parkinson’s Disease (Table 1).

**Table 1.**
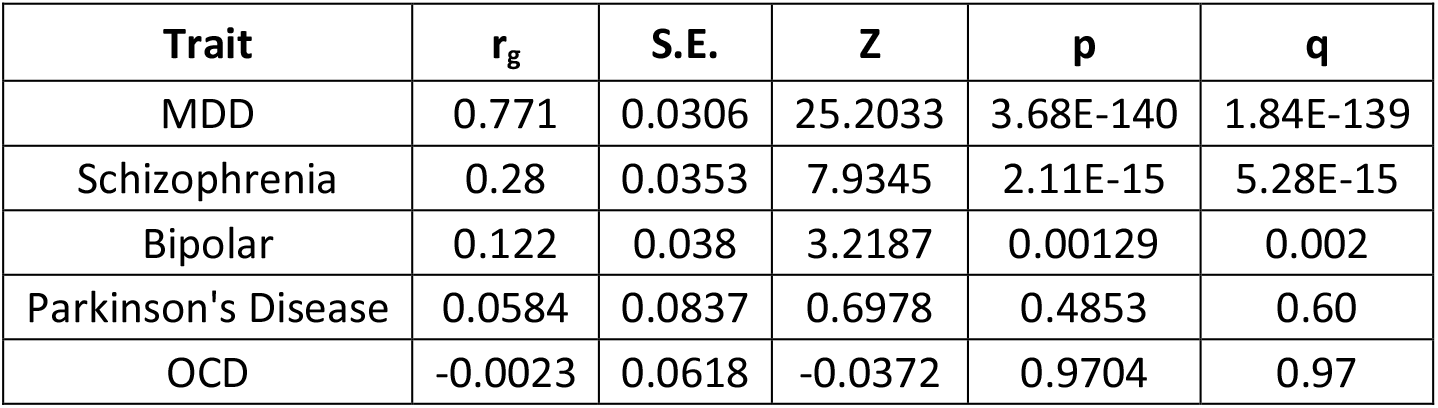
Genetic correlations of mood instability with psychiatric phenotypes. rg = genetic correlation with mood instability, S.E. = standard error of the genetic correlation, Z = the test statistic, p = the p value, q the False discovery rate corrected p value. MDD = major depressive disorder, PTSD = post-traumatic stress disorder.

### PRS and brain MRI structural and functional outcomes

Associations between PRS for anhedonia and total brain tissue volumes are presented in Table S3. Greater polygenic risk for anhedonia was associated with lower total grey matter volume, but not with total white matter volume or with total ventricular CSF volume.

Associations were then assessed between PRS for anhedonia and volumes of 15 cortical and subcortical regions of interest (ROIs were derived *a priori* from meta-analysis and literature review; please see Table S4 and Supplementary Methods). Greater genetic risk score for anhedonia was associated with smaller volumes for nucleus accumbens, insular cortex, medial frontal cortex, orbitofrontal cortex, middle frontal gyrus and anterior temporal fusiform cortex. No significant associations were observed for the remaining nine ROIs. The locations of the ROIs showing significant association with the PRS are displayed in Supplementary Figure S12, and a point-range plot showing the association evidence for ROI volumes is displayed in Figure 2A.

**Figure 2.**
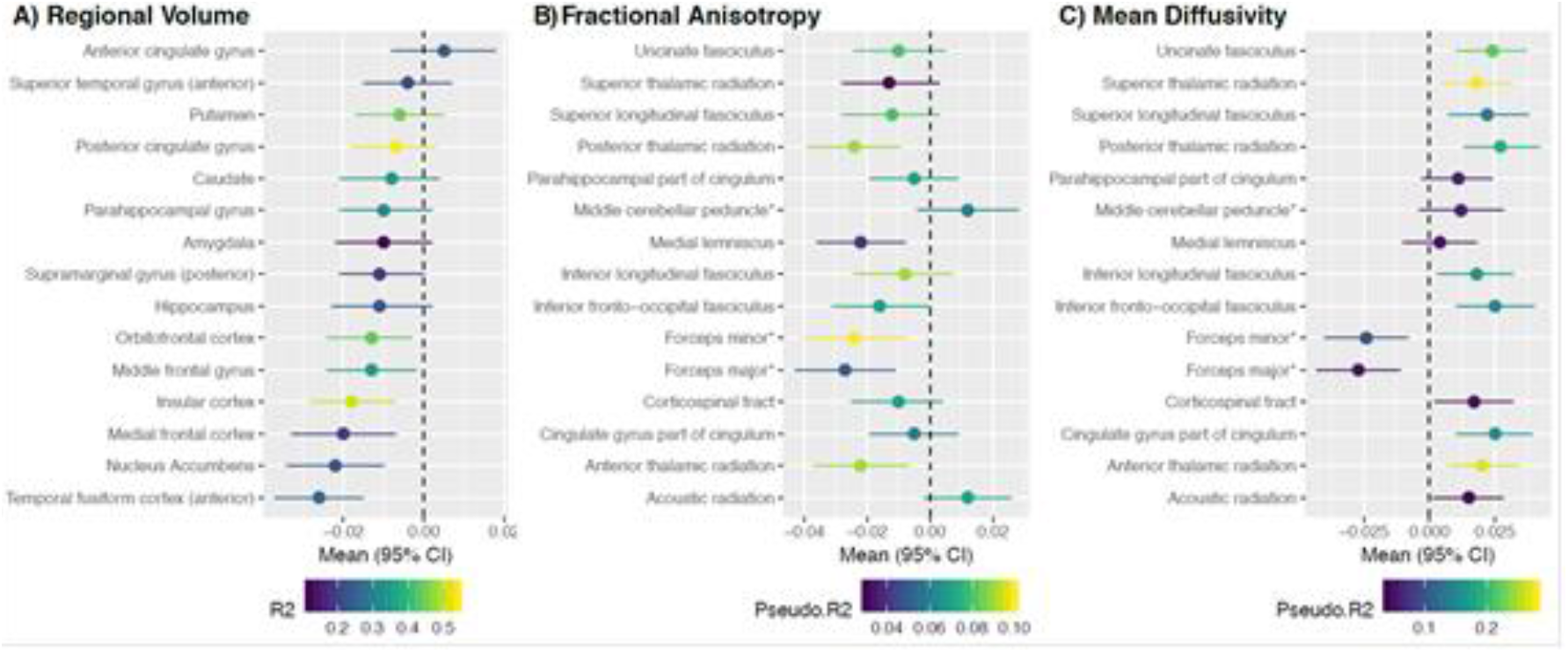
Plots of associations (regression coefficients and 95% confidence intervals (CI)) between polygenic risk for anhedonia and A) regional volumes of cortical/subcortical ROIs; B) tract-specific fractional anisotropy; and C) tract-specific mean diffusivity.

In subsequent analyses of association with white matter integrity, greater PRS for anhedonia was associated with a higher mean diffusivity (MD) (Table S5) but no association was observed between PRS for anhedonia and fractional anisotropy (FA). PRS associations were assessed with 15 individual white matter tracts (bilateral tracts were combined in the same models and models were adjusted for hemisphere; see Supplementary Methods, Supplementary Table S6 and S7). Higher PRS for anhedonia was associated with lower FA in five of the fifteen tracts (anterior thalamic radiation, forceps major, forceps minor, medial lemniscus, posterior thalamic radiation) (Figure 2B). Higher PRS for anhedonia was also associated with higher MD in 12 of the 15 tracts (all apart from forceps major, medial lemniscus and parahippocampal part of cingulum) (Figure 2C). There was no association between PRS for anhedonia and functional MRI activity during the emotion-processing task (median Blood Oxygen Level Dependent signal for the face vs. shape contrast) (Table S8).

## Discussion

These analyses represent the largest genetic association study of anhedonia performed to date. We identified eleven genetic loci associated with anhedonia in the UK general population. Within each associated region there were a number of genes that could have a functional impact on anhedonia and the pleasure cycle (considered in detail below). Consistent with an RDoC approach focusing on transdiagnostic traits, we found strong genetic correlations between anhedonia and MDD, schizophrenia and bipolar disorder, but not between anhedonia and OCD or Parkinson’s Disease. Despite the lack of correlation with Parkinson’s Disease, several of the loci identified in our GWAS include genes with known association with Parkinson’s Disease (see below). We also report the first investigation of associations between genetic loading for anhedonia and both brain structure and brain function.

### Genes within anhedonia-associated loci

Within the chromosome 1 locus there are multiple *RGS* genes, most notably *RGS1* and *RGS2*, encoding regulators of G-protein signalling that show prominent expression in the brain^33^. *RGS2* has previously been identified as a modulator of *LRRK2*^34^ expression, a gene known to be a genetic cause of Parkinson’s Disease. *RGS2* has also been associated with symptom severity in schizophrenia^35^ and lower expression of *RGS2* may be related to depression-like behaviours in animal models^36^.

*EPHB1* on chromosome 3 encodes an ephrin receptor tyrosine kinase identified in a GWAS of antidepressant response^37^, and is associated with symptoms of schizophrenia in Chinese Han populations^38^, and with susceptibility to Parkinson’s Disease^39^.

At the more centromeric locus on chromosome 11, *GRM5* encodes a metabotropic glutamate receptor that has been extensively studied in relation to MDD^40, 41^ and schizophrenia^42, 43^. Assessments in mice have also found that agonists of *GRM5* attenuate Parkinsonian motor deficits via striatal dopamine depletion^44^. The Genotype Tissue Expression (GTEx) database^33^ shows prominent expression of *GRM5* in the brain, especially within the nucleus accumbens, a region with a major role in the prediction of reward^45, 46^. As discussed further below, it is of note that we found that greater PRS for anhedonia was associated with smaller volume of the nucleus accumbens.

Another likely candidate gene at this chromosome 11 locus is *DISC1FP1*. *DISC1* (Disrupted in Schizophrenia 1), a gene located on chromosome 1, is part of a chromosome 1:11 translocation that increases risk of schizophrenia, schizoaffective disorder and bipolar disorder^47^. *DISC1FP1* is disrupted by this translocation, impacting on both intracellular NADH oxidoreductase activities and protein translation^48^.

At the more telomeric locus on chromosome 11, *NCAM1* (neural cell adhesion molecule 1) has been implicated as a potential link between depressive symptoms and brain structure^49^, specifically decreased FA. *NCAM1* has been reported to be present at increased levels in the amygdala^33^, another brain region associated with the pleasure cycle, in depressed subjects^50^. However, we did not find any association between PRS for anhedonia and either amygdala volume or functional activity in the amygdala during an emotion processing task. It is possible that *NCAM1* may exert its effects via expression in other brain regions.

Another gene of interest at this second chromosome 11 locus is *DRD2*, encoding the dopamine D2 receptor. There is an extensive literature on the importance of *DRD2* for the psychopharmacology of both schizophrenia and MDD^51,52^.

The chromosome 12 locus contains only *LOC* and *LINC* genes. These encode different classes of functional RNA but little is known about their function beyond possible post-transcriptional regulation of other gene products.

The more centromeric locus on chromosome 14 contains *PRKD1*, which encodes a serine/threonine-protein kinase identified in a GWAS of schizophrenia^16^. The more telomeric locus on chromosome 14 contains the gene *SLC8A3*, a gene involved in maintaining Ca^2+^ homeostasis within a variety of tissues, including neurons. *SLC8A3* may also play a role in Parkinson’s Disease^53^.

*ISLR2* on chromosome 15 is involved in neurodevelopment^54^ but its role in psychiatric and neurological traits is not well characterised. Another candidate gene within this locus, *NRG4*, encodes a neuregulin protein that activates type-1 growth factor receptors. Recent work has shown that NRG4 acts a regulator of the growth and elaboration of pyramidal neuron dendrites in the developing neocortex^55^. Pyramidal neurons have been directly implicated in the pathophysiology of schizophrenia^56^.

Finally, on chromosome 18, *DCC* (the most significant hit) encodes a netrin 1 receptor with a role in axon guidance, and has been previously reported to be associated with anhedonic phenotypes in mice and humans^57^ and potentially with schizophrenia pathogenesis^58^. We have previously identified *DCC* in GWAS of mood instability^59^, suicidality^60^ and multisite chronic pain^61^.

### Brain Structure and function

Our findings on the relationship between genetic loading for anhedonia and brain structure and function are of considerable interest. Greater levels of anhedonia, in healthy and clinical populations, have been linked to altered functional activity (and less consistently to reduced volume) in frontal/striatal regions involved in reward or pleasure processing^10–12, 62–66^. The regions most consistently implicated include nucleus accumbens, caudate, putamen, medial frontal cortex and orbitofrontal cortex.

We found that increased genetic risk for anhedonia was associated with smaller volumes of nucleus accumbens, medial frontal cortex and orbitofrontal cortex (Figure 2A; Supplementary Figure 1). Nucleus accumbens is involved in reward value and pleasure processing^64, 67^. Orbitofrontal cortex is involved in representation of reward value and social reward dependence^65, 68^, and medial frontal cortex has been linked to pleasure processing^64^. These cortical/subcortical volume findings are therefore consistent with an association between genetic risk for anhedonia and abnormal reward/pleasure processing^67^.

Increased genetic risk for anhedonia was also associated with smaller volumes of insular cortex (associated with emotion processing^69^) and fusiform cortex. Most ROIs in our analysis were selected on the basis that they were associated with smaller volumes in MDD versus healthy controls in the most recent MDD brain structure meta-analysis^70^. Our findings are consistent with two scenarios: either genes for anhedonia partly mediate the association between the ROI volumes and MDD, or these genes exhibit horizontal pleiotropy and affect both phenotypes via separate mechanisms. Consistent with our finding of widespread associations of genetic risk for anhedonia with cortical/subcortical volumes, we also found an association with reduced total grey matter volume, adjusted for head size.

Evidence of reduced white matter integrity (FA) in several tracts in individuals scoring higher on measures of anhedonia has been reported^14, 71^. We found that higher values for the anhedonia PRS were associated with higher MD (reflecting poorer white matter integrity), both in a general factor, and in most individual tracts. Several tracts also showed reduced FA.

We did not, however, find any association between PRS for anhedonia and functional brain activity. UK Biobank ROIs were selected based on average responses during an emotion processing task; it is therefore plausible that effects would emerge with the use of a reward processing task, or by applying whole-brain voxel-wise analyses, to include further reward/pleasure processing regions.

### Strengths and limitations

This study is the largest GWAS of anhedonia to date, and substantially contributes new knowledge on the biology of this important transdiagnostic symptom. We conducted analyses within a large population-based cohort. Our primary analysis included individuals with mental health histories, but these represent only a small proportion of a total sample, which is an order of magnitude larger than any previous study of this kind. Subclinical anhedonia is common, and associated with increased risk of later mental illness^72^. This idea is reinforced by the significant genetic correlations with other psychiatric disorders. Additionally, we used a multi-level ordinal phenotype, resulting in more power to detect associations than with the more common dichotomised (‘non-anhedonic’ versus ‘anhedonic’) analyses^73^. Identification of loci associated with population-level anhedonia may be important from a precision medicine perspective, for example in terms of developing stratified medicine approaches to identify individuals at high-risk of developing psychiatric disorders.

Notably, however, the UK Biobank cohort has a degree of selection bias. In general, volunteers are typically healthier and of higher socioeconomic status and education level than the general population^74^, which may have led to *under*-representation of predisposing alleles in the higher levels of the anhedonia outcome variable. This bias may have been further amplified in the sub-sample who attended the imaging visit (these participants also showed somewhat reduced likelihood of reporting recent symptoms of anhedonia). Therefore, it is plausible that the strength of reported associations may be an underestimate of the true population value, and there may also be an inflated type 2 error rate.

The measure of anhedonia employed here was a question from a depression screening instrument, the PHQ-9, assessing frequency of anhedonia within the past two weeks. This item therefore measures state anhedonia at a single time point, and could be influenced by environmental factors, such as season or current health status^75, 76^. However, we have assumed here that despite transient environmental and physical factors, individuals who are prone to trait anhedonia will be more likely to report higher frequency of anhedonia at any given time point, and it seems likely that a recent anhedonia phenotype will be enriched for individuals with stronger genetic predisposition compared to a similar phenotype based on lifetime incidence. In line with this assumption, the anhedonia item of the PHQ-9 at a single time point has shown utility in predicting longitudinal brain structural change^77^.

The GWAS treated the ordinal responses to the frequency of anhedonia question as linear. This approach is likely to have had a minimal impact on the BOLT-LMM GWAS results. Although the distance between points on the anhedonia scale is not evenly spaced, each point can reasonably be considered to be higher than the point before it.

The anhedonia measure did not enable determination of specific anhedonia subtypes. Existing validated instruments typically divide anhedonia into physical and social subscales^19^, or into anticipatory versus consummatory components of pleasure^78^. Future studies using more detailed anhedonia scales may be of use in examining the extent of genetic overlap between anhedonia subtypes.

## Conclusion

We report the largest GWAS to date of anhedonia, a common symptom associated with several psychiatric disorders. We identified 11 novel genetic loci and our findings indicate substantial genetic overlap between anhedonia and several psychiatric disorders including MDD, schizophrenia and bipolar disorder. PRS analyses revealed association between genetic loading for anhedonia and smaller volumes of several brain regions, and poorer white matter integrity. Taken together, these findings provide important insights into the neurobiology of an important but under-studied psychiatric symptom and strongly support the proposition that genetic predisposition to anhedonia may influence brain structure and function.

## URLs

UK Biobank genetic data release information - https://data.bris.ac.uk/datasets/3074krb6t2frj29yh2b03x3wxj/UK%20Biobank%20Genetic%20Data_MRC%20IEU%20Quality%20Control%20version%201.pdf

## Acknowledgements

JW is supported by the JMAS Sim Fellowship for depression research from the Royal College of Physicians of Edinburgh (173558). AF is supported by an MRC Doctoral Training Programme Studentship at the University of Glasgow (MR/K501335/1). RJS is supported by a UKRI Innovation-HDR-UK Fellowship (MR/S003061/1). KJAJ is supported by an MRC Doctoral Training Programme Studentship at the Universities of Glasgow and Edinburgh. DJS acknowledges the support of a Lister Prize Fellowship (173096) and the MRC Mental Health Data Pathfinder Award (MC_PC_17217). RAIB is supported by a British Academy Post-Doctoral Fellowship.

