## Supplementary figures and images for "Novel genome-wide associations for anhedonia, genetic correlation with psychiatric disorders, and polygenic association with brain structure"

### Figure 2

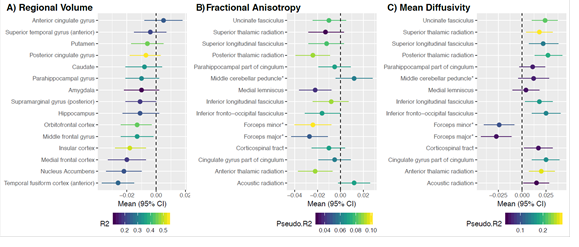

### Figure S1

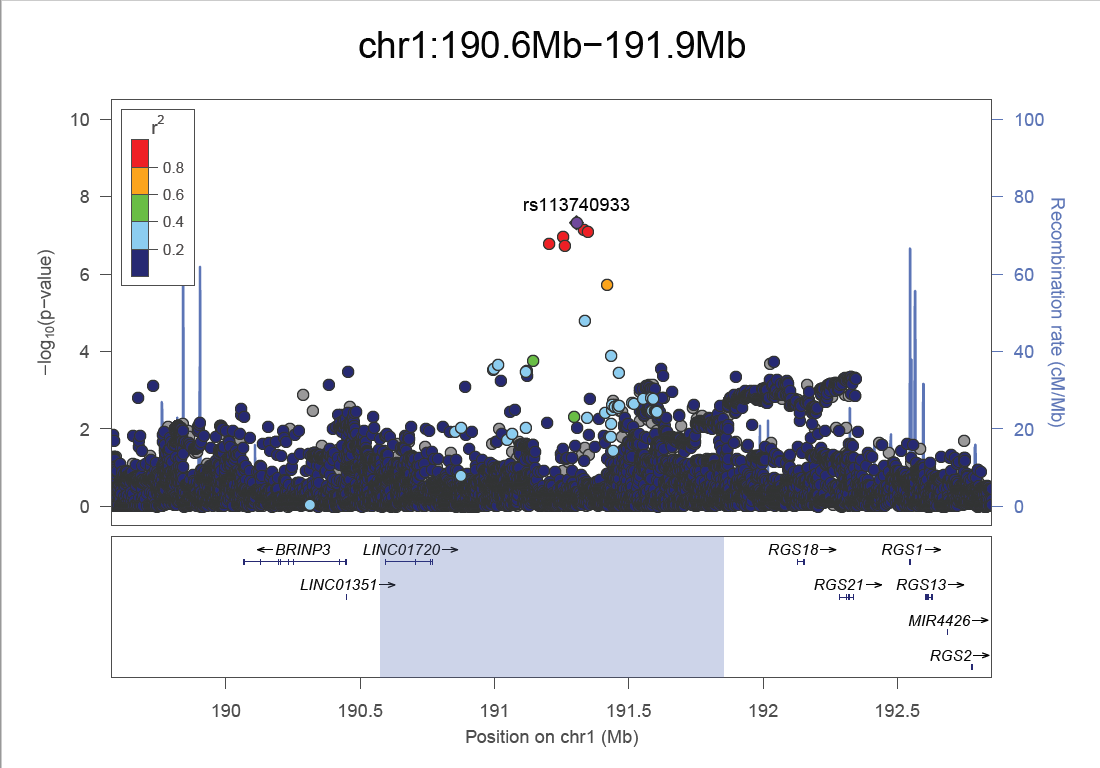

### Figure S2

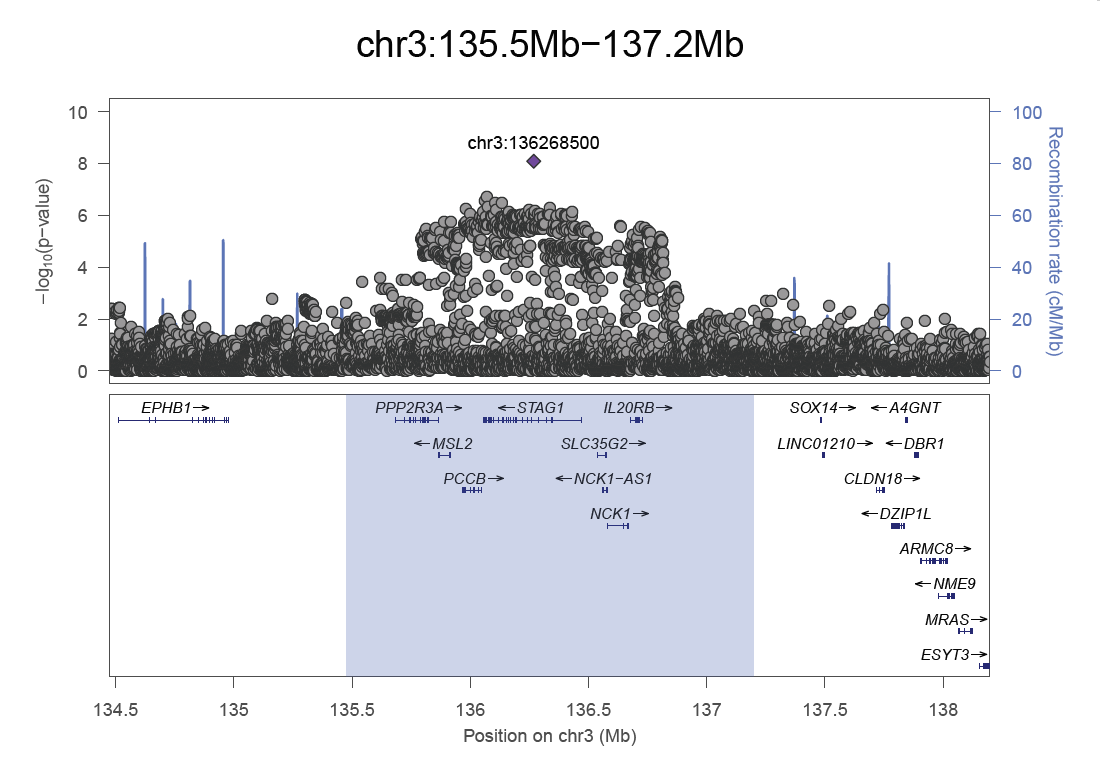

### Figure S3

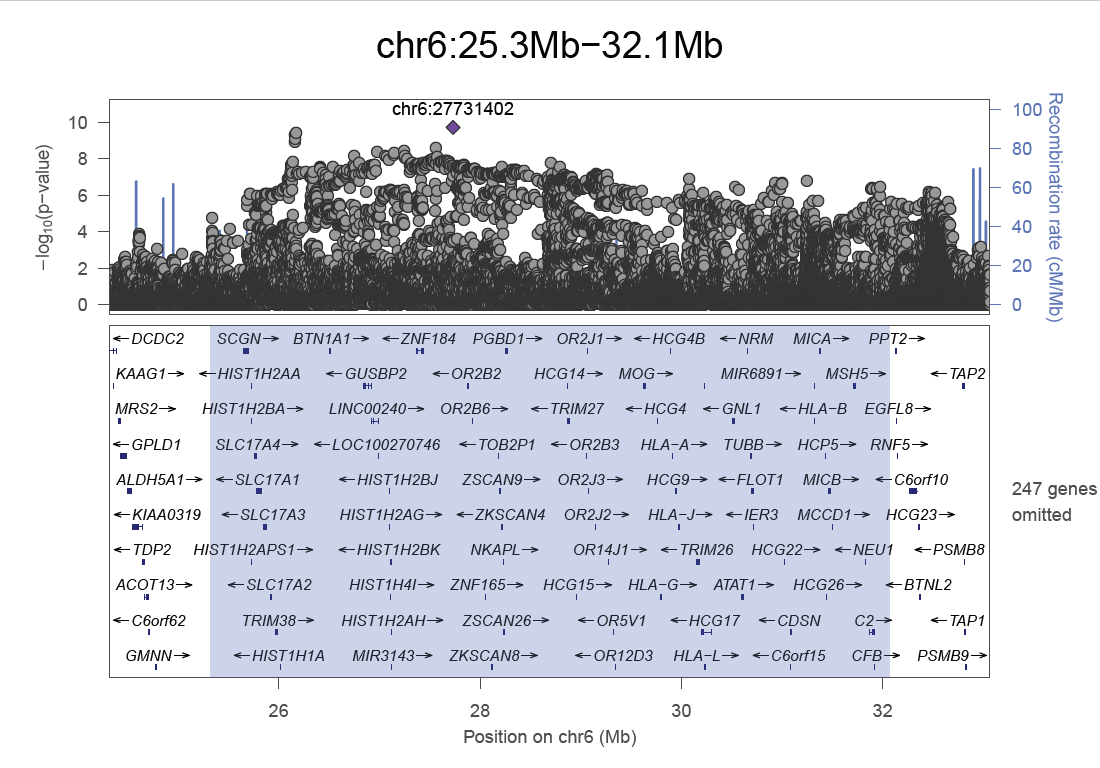

### Figure S4

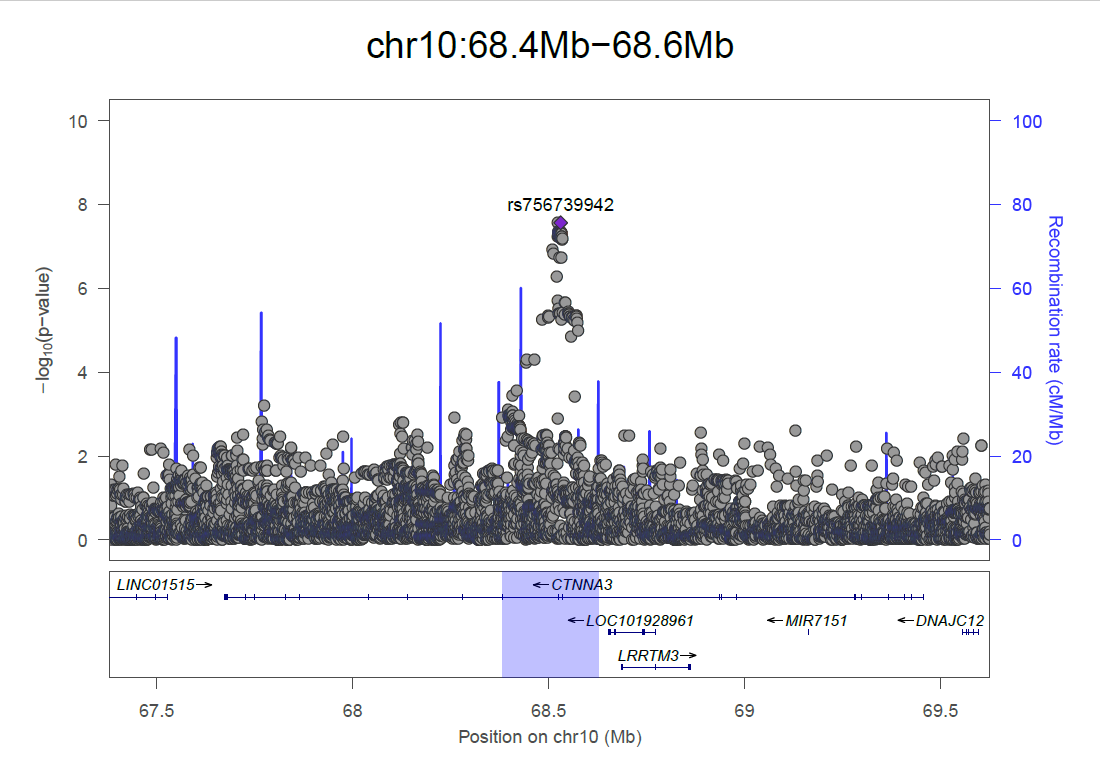

### Figure S5

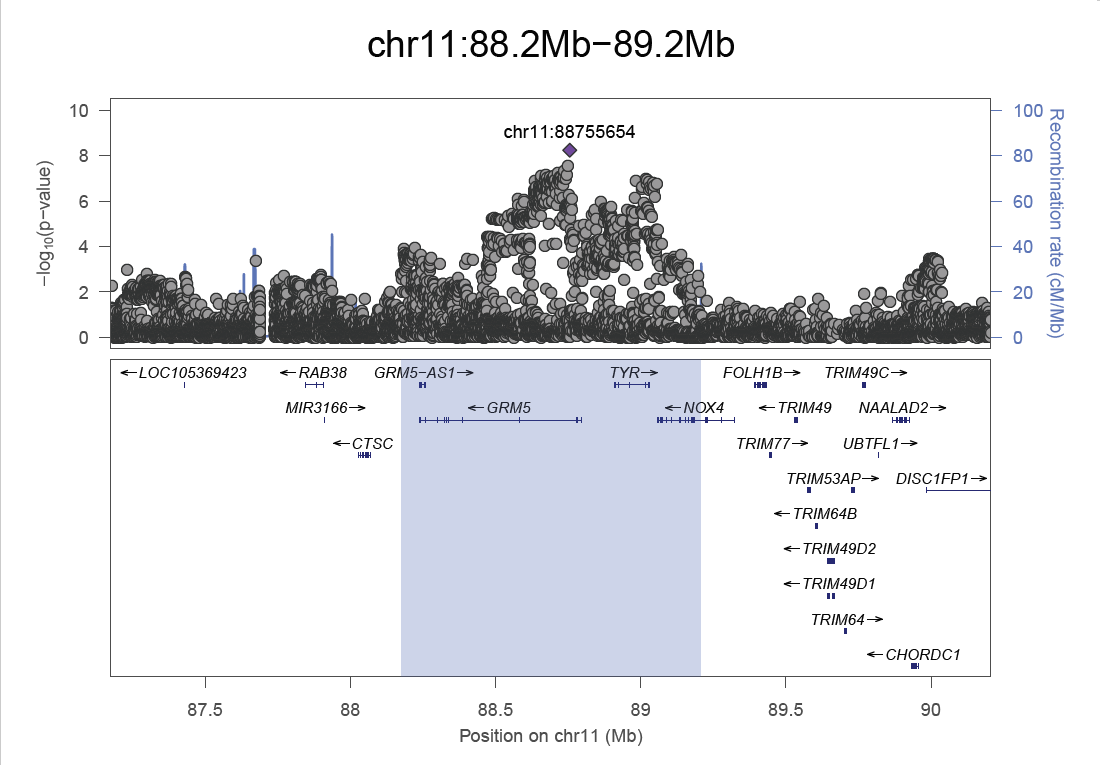

### Figure S6

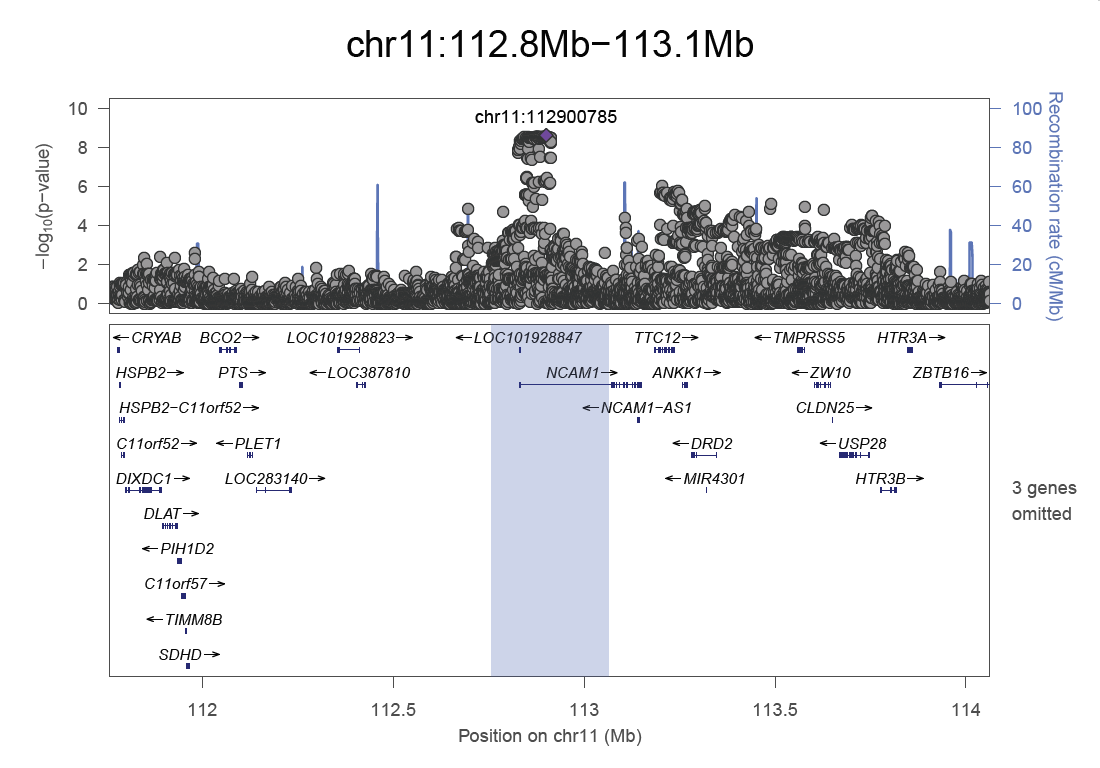

### Figure S7

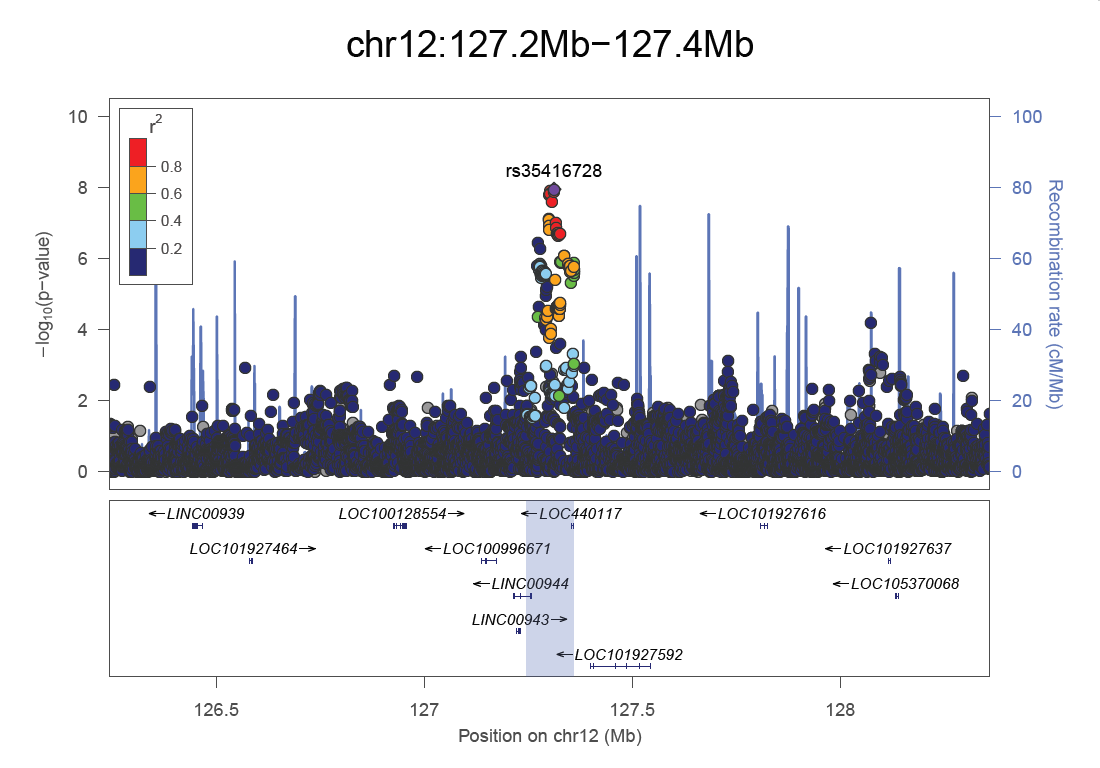

### Figure S8

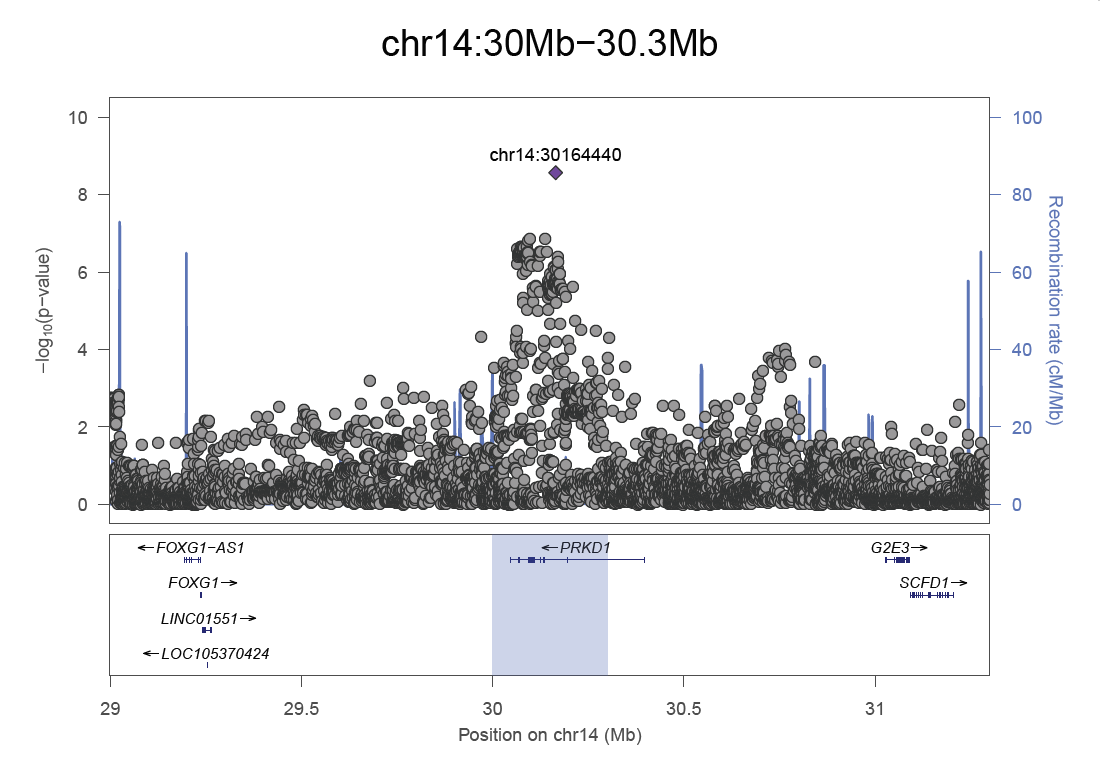

### Figure S9

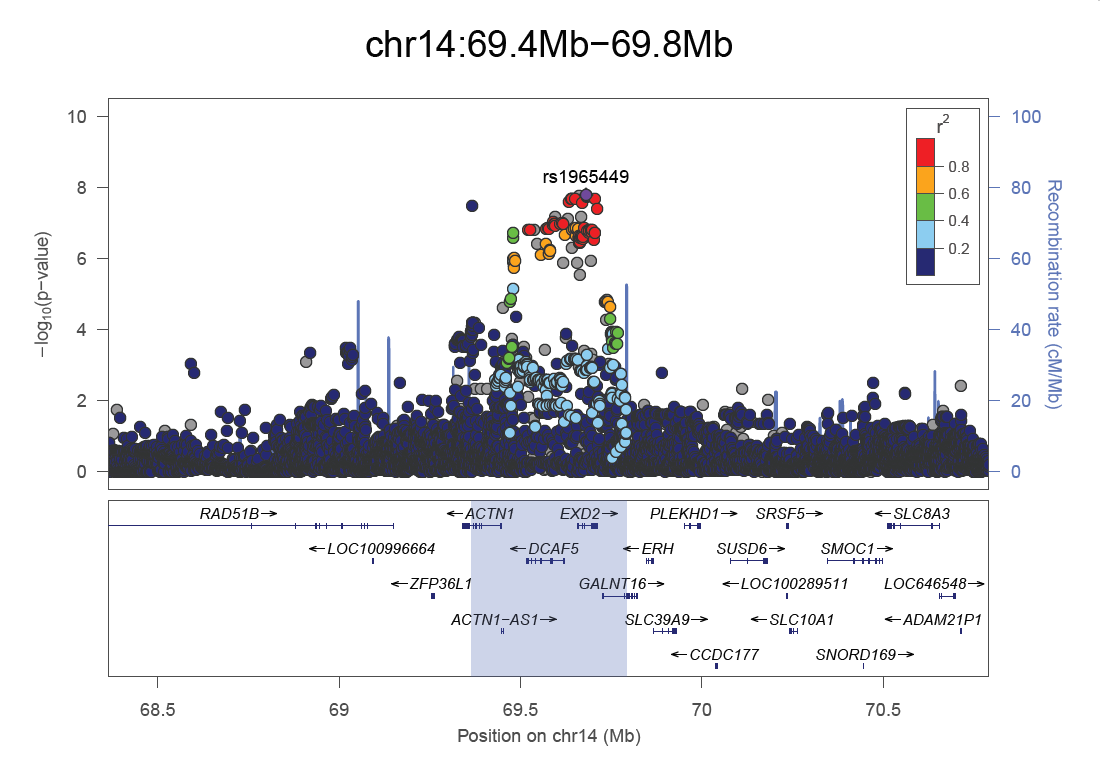

### Figure S10

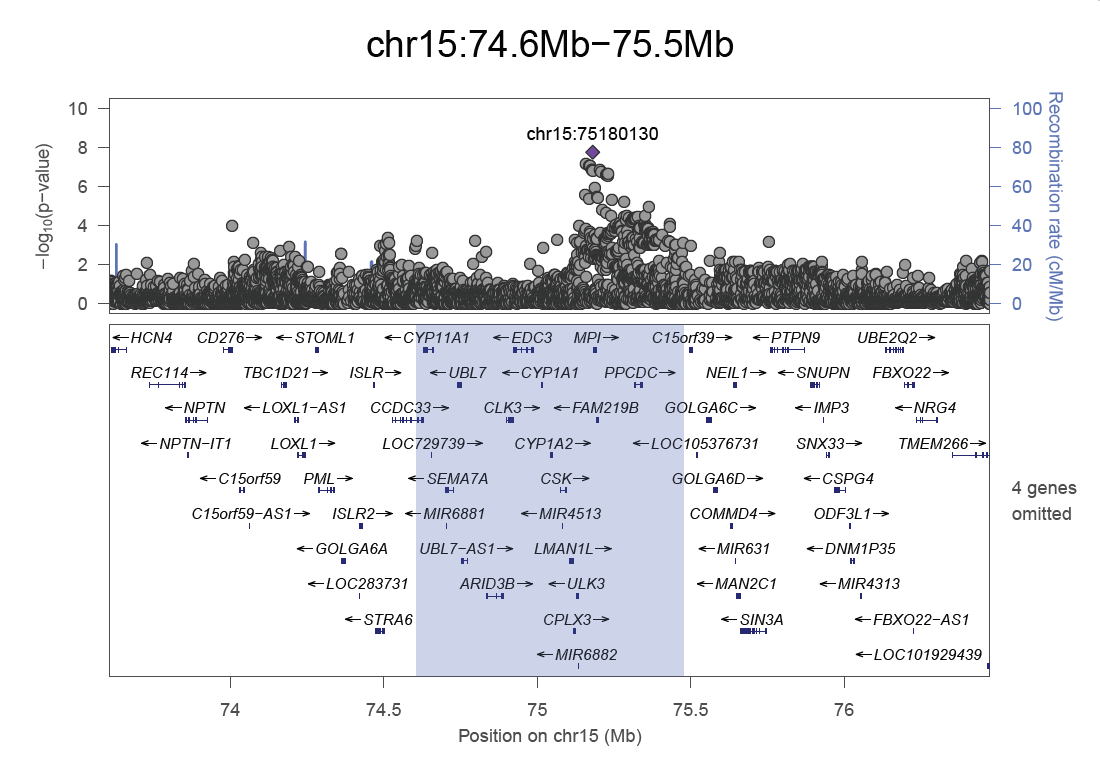

### Figure S11

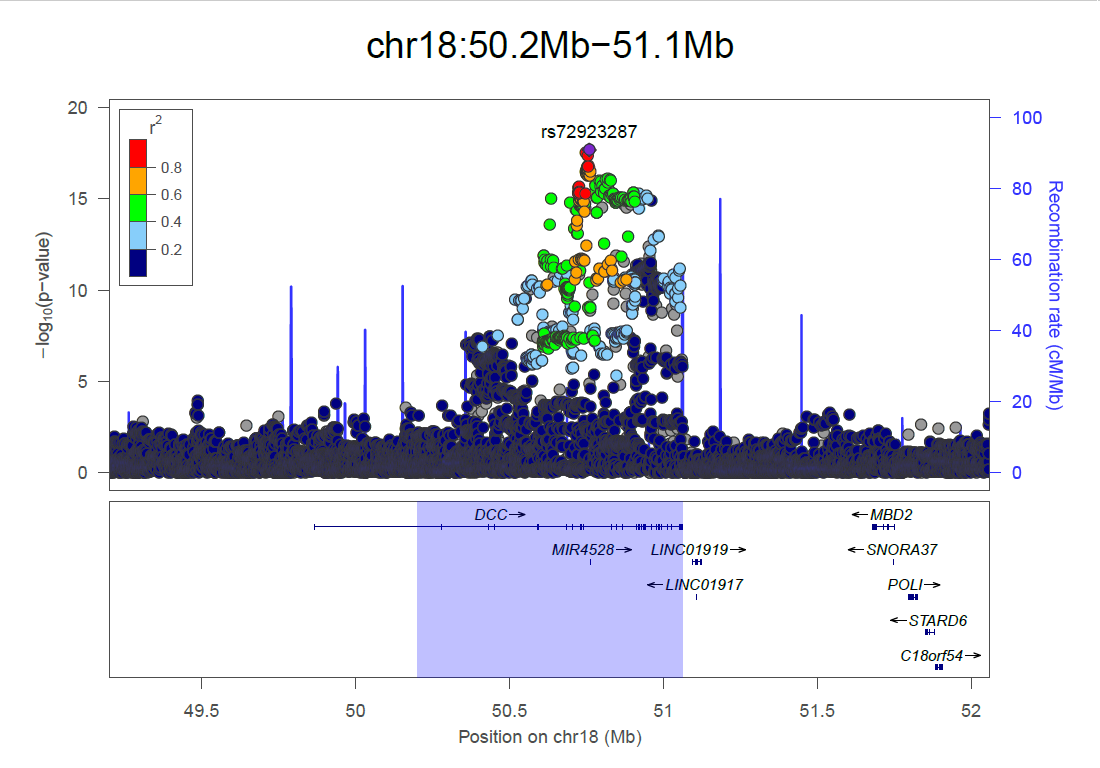

### Figure S12

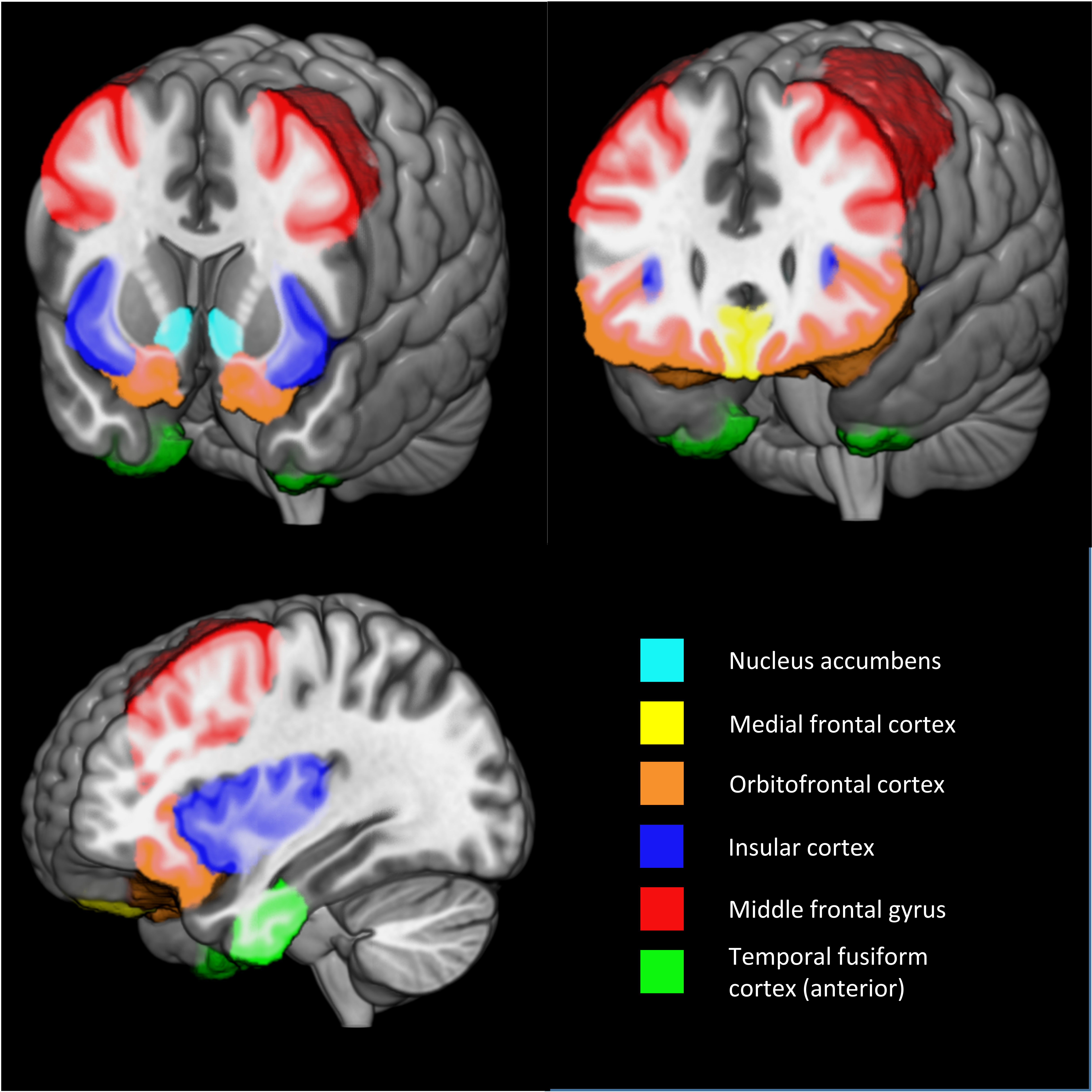
